# Inhibition of 15-PGDH Protects Mice from Immune-mediated Bone Marrow Failure

**DOI:** 10.1101/2020.04.07.030312

**Authors:** Julianne N.P. Smith, Folashade Otegbeye, Alvin P. Jogasuria, Kelsey F. Christo, Stanton L. Gerson, Sanford Markowitz, Amar B. Desai

## Abstract

Aplastic anemia (AA) is a human immune mediated bone-marrow failure syndrome that is treated by stem cell transplantation for patients who have a matched related donor or immunosuppressive therapy (IST) for those who do not. Responses to IST are variable, with patients still at risk for prolonged neutropenia, transfusion-dependence, immune suppression, and severe opportunistic infections. Therefore, additional therapies to accelerate hematologic recovery in patients receiving front line IST are needed. We have shown that inhibiting 15-hydroxyprostaglandin dehydrogenase (15-PGDH) with the small molecule SW033291 (PGDHi) increases bone marrow (BM) prostaglandin E2 levels, expands hematopoietic stem cell (HSC) numbers, and accelerates hematologic reconstitution following murine BM transplantation. We now report that in a murine model of immune-mediated BM failure, PGDHi therapy mitigated cytopenias, increased BM HSC and progenitor cells, and significantly extended survival compared to vehicle-treated mice. PGDHi protection was not immune-mediated, as serum IFNγ levels and BM CD8+ T lymphocyte frequencies were not impacted. Moreover, dual administration of PGDHi plus low dose IST enhanced total white blood cell, neutrophil and platelet recovery, achieving responses similar to maximal dose IST at lesser toxicity. Together these data demonstrate that PGDHi can complement IST to accelerate hematologic recovery and reduce morbidity in severe AA.

## Introduction

Acquired aplastic anemia (AA) is a primary bone marrow (BM) failure syndrome in which hematopoietic stem cells (HSC) are depleted in an immune-mediated fashion (1, 2). AA is rare, with an estimated 900 new cases per year in the U.S., and onset is typically in childhood or adolescence (3). AA is fatal if untreated and poses a high burden of frequent, long-term transfusions and associated complications. Treatment options are currently limited to allogeneic bone marrow transplantation for young patients with a matched related donor, or immunosuppressive therapy (IST) using cyclosporine (CsA) and horse antithymocyte globulin as a first-line therapy for those who do not (4). Due to the potential for severe hepato- and nephrotoxicity, however, CsA dosing often requires adjustment to achieve maximal efficacy at the lowest possible dose (5). Additional agents including G-CSF and more recently, eltrombopag, are used to augment hematologic recovery in AA patients not responding well to IST. These agents are inconsistently effective (6), and G-CSF has been associated with clonal evolution in some studies (7). Therefore, there is a great clinical need for additional well-tolerated therapeutic strategies to complement and further extend the efficacy of first-line therapies.

AA pathogenesis involves the activation and proliferation of autoreactive T cells, which home to the BM, resulting in direct and microenvironmentally-mediated HSC depletion (8). HSCs are likely depleted via both apoptosis and functional exhaustion (9–11). Prostaglandin E2 (PGE2) enhances stem cell survival and stem and early progenitor cell proliferation (12–14). Moreover, we have previously shown that inhibition of the PGE2-degrading enzyme 15-hydroxyprostaglandin dehydrogenase (15-PGDH) increases BM PGE2 levels and increases HSCs and hematologic recovery following transplantation (15, 16). Here we sought to test the capacity for 15-PGDH inhibition (PGDHi) to ameliorate disease alone and in combination with IST in a murine model of immune-mediated BM failure. Our data demonstrate that PGDHi limits pancytopenia, BM aplasia, and HSC loss, but does not impact inflammation and T lymphocyte frequencies in the BM. Further, PGDHi augments the efficacy of low dose cyclosporine, demonstrating the potential clinical utility of PGDHi to augment therapeutic effect in AA patients who are dose-limited due to cyclosporine-associated toxicity or who are refractory to standard doses of immunosuppressive therapy.

## Results

### 15-PGDH inhibition ameliorates cytopenias, extends survival, and preserves hematopoietic progenitors in murine aplastic anemia

In order to test the therapeutic utility of PGDHi during severe aplastic anemia (AA) we modeled AA in mice by infusing sublethally-irradiated recipient C57BL/6 x BALB/c F1 hybrid mice (H^b/d^) with histocompatibility-mismatched lymphocytes from C57BL/6 (H^b/b^) donors. This well-established model results in immune-mediated marrow failure and elicits little inflammation in non-hematopoietic organs (17). We tested effects of PGDHi using (+)-SW033291, a potent and specific small molecule 15-PGDH inhibitor (15, 16). To specifically determine the impact of PGDHi on established disease, rather than on the establishment of immune-mediated marrow failure, the 15-PGDH inhibitor (+)SW033291 was administered beginning 4 days post-disease induction (**Fig. 1A**). 13 days post-disease induction, vehicle-treated mice are severely leukopenic, neutropenic, and thrombocytopenic relative to mice that received radiation alone. PGDHi treatment resulted in significant increases in white cells, neutrophils and platelets (**Fig. 1B**), though had no impact on red blood cell counts (**Supp. Fig. 1**). Thus 15-PGDH inhibition prevented the severe pancytopenia of this autoimmune bone marrow failure model. This improvement was further reflected in improved survival of PGDHi treated mice. Whereas AA was uniformly lethal in vehicle-treated mice, PGDHi-treated mice showed significantly extended overall survival time with long term protection in 10% of animals (**Fig. 1C**).

**Figure 1:**
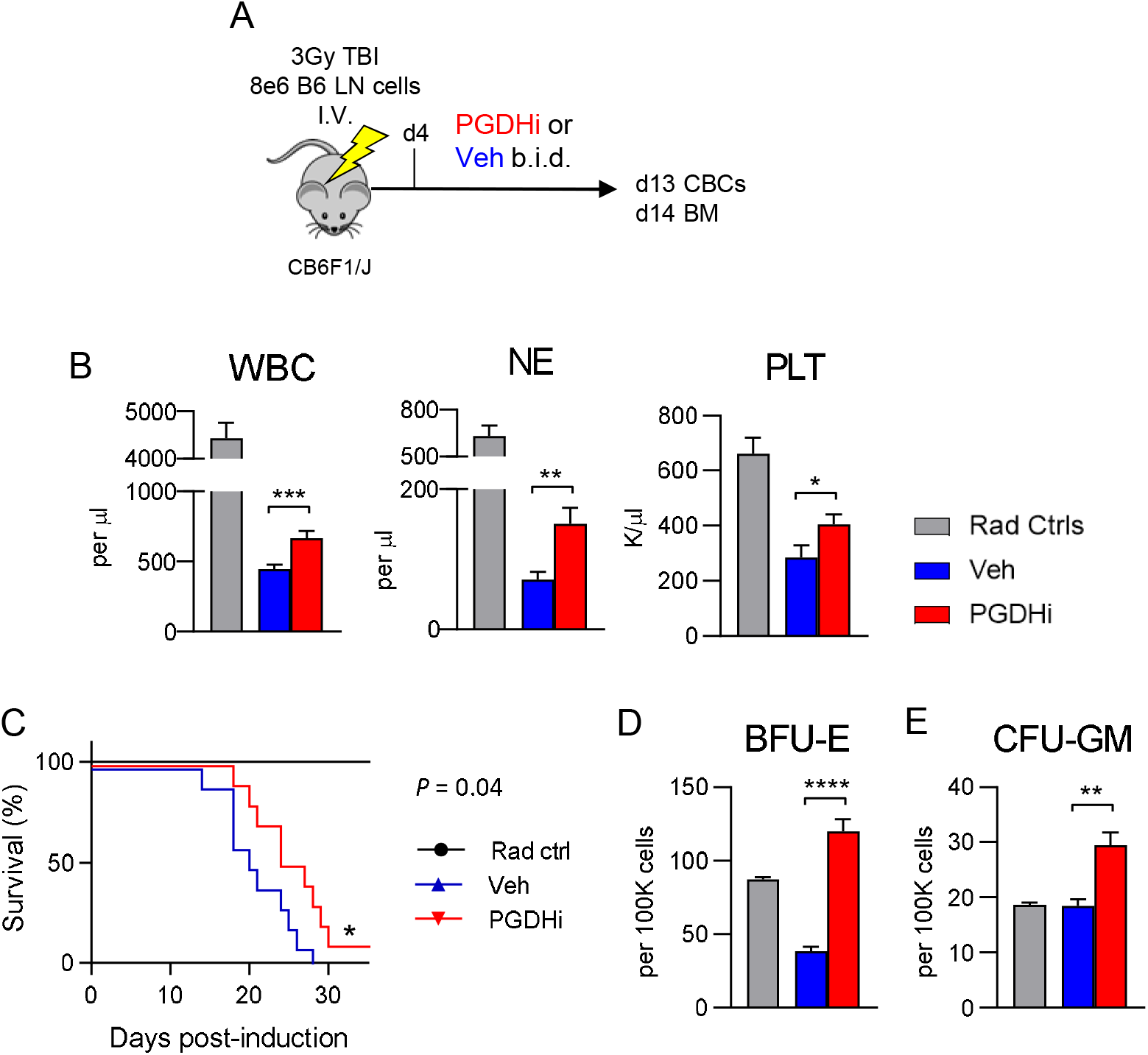
15-PGDH inhibition attenuates the severity of murine autoimmune bone marrow failure. (**A**) Schematic depicting the disease model, in which CB6F1/J mice are induced via 3Gy irradiation and infusion with 8e6 C57BL/6 lymph node (LN) cells, and subsequently treated with PGDHi or vehicle (Veh) control, beginning at day 4. (**B**) Complete blood count analysis of white blood cells (WBCs), neutrophils (NE), and platelets (PLT) in radiation alone controls (grey bars), and aplastic anemia mice treated with vehicle- (blue), or PGDHi- (red), 13 days post-induction. N=21 experimental mice/group. (**C**) Kaplan-Meier survival curve. N=5 Rad ctrl, and 10 Veh- and PGDHi-treated mice/group. (**D-E**) Quantification of burst-forming unit erythroid (BFU-E) and colony-forming unit granulocyte, monocyte (CFU-GM) per 100K BM cells. N=5-6 mice/group.

To determine if PGDHi may mitigate pancytopenia via actions on hematopoietic progenitor cells, we measured colony-forming capacity in the BM of AA mice 14 days post-induction. Whereas burst forming units-erythroid (BFU-E) were severely diminished in vehicle-treated AA mice, PGDHi-treated mice demonstrated a 3- and 1.5-fold increase in BFU-E and colony-forming units-granulocyte/macrophage (CFU-GM), respectively (**Fig. 1D-E**).

### PGDHi protection is not immune-mediated

Interferon gamma (IFNγ) and TNFα are cytokines classically associated with AA pathogenesis and are known to exhaust and deplete HSCs (1, 18, 19). To determine if PGDHi limits AA severity by attenuating inflammation, we measured circulating levels of the disease-driving cytokines IFNγ and TNFα, as well as IL-6 and IL-1β. With the exception of IL-1β, these factors were upregulated at day 14 post-induction compared to radiation controls, however, levels were not impacted by PGDHi treatment (**Fig. 2A**). Analysis of an extended panel of cytokines and chemokines confirmed the general lack of impact of PGDHi on these effectors (**Supp. Fig. 2**). We next quantified BM lymphocytes to address the potential for PGDHi to impair T cell homing to or proliferation in the local microenvironment. Although we observed a PGDHi-dependent trend towards reduced total CD90^+^ cells, the absolute numbers and frequencies of CD8^+^ and CD4^+^ T lymphocytes were unchanged between PGDHi- and vehicle-treated mice (**Fig. 2B-D**). Additionally, the expression levels of CD90, FasL, and CD11a on CD8^+^ (**Supp. Fig. 3A**) and CD4^+^ (**Supp. Fig. 3B**) T cells were not impacted by PGDHi. Together these data support that PGDHi does not limit AA severity through an immune-modulatory mechanism, thus raising the possibility that PGDHi therapy could be synergistic with immunosuppressive therapy (IST) as treatment for AA.

**Figure 2:**
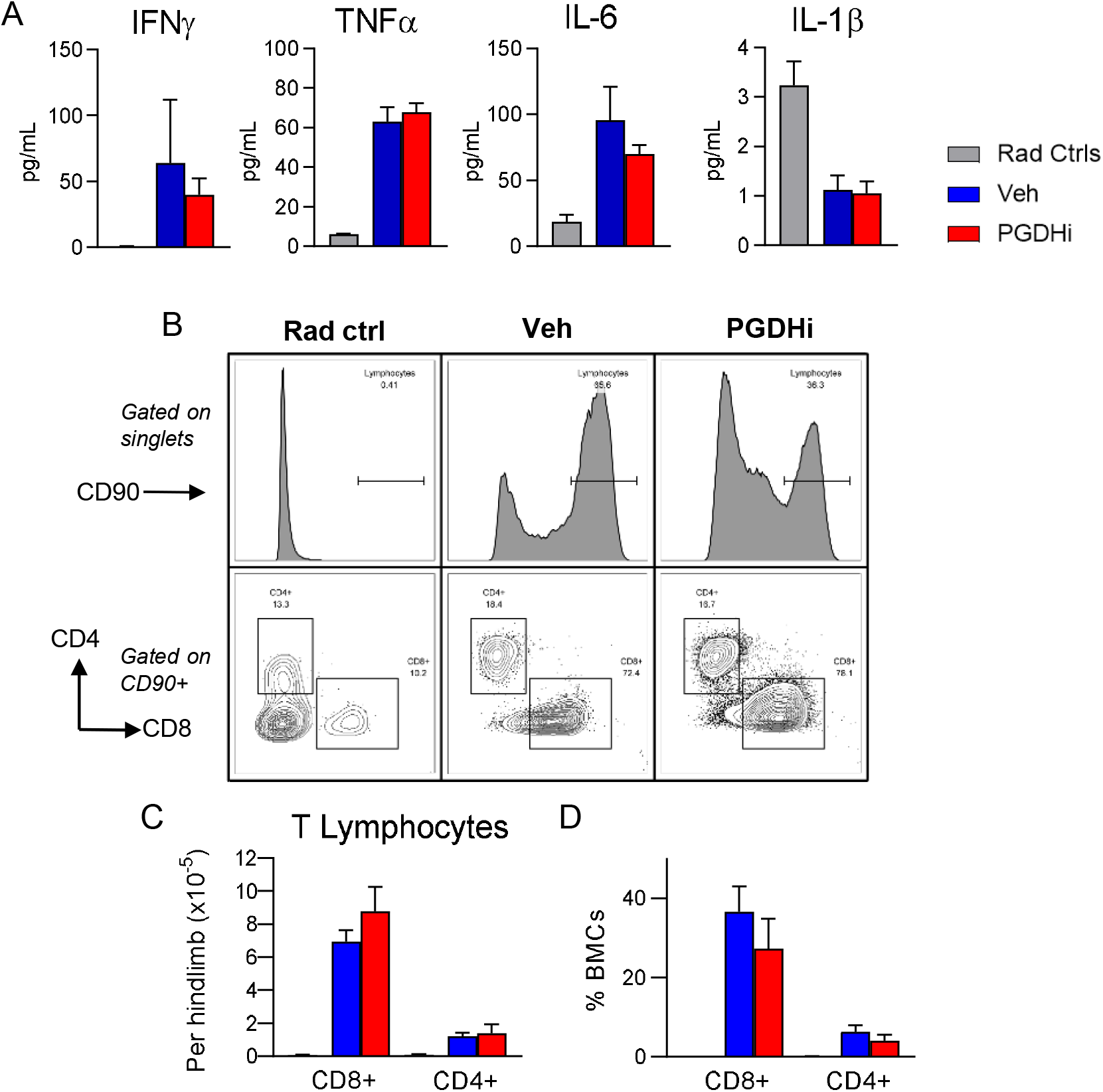
PGDHi does not alter inflammatory cytokine levels in peripheral blood serum or BM lymphocyte frequencies. (**A**) Measurement of IFNγ, TNFα, IL-6, and IL-1β in peripheral blood serum of radiation control (grey), and aplastic anemia mice treated with vehicle- (blue), and PGDHi- (red), 14 days post-induction. N=17-18 experimental mice/group. (**B**) Representative CD90 histograms (top; gated on single bone marrow cells), and CD8 (x) and CD4 (y) plots (bottom; gated on CD90+ bone marrow cells) from radiation control (Rad ctrl), and Veh-, and PGDHi-treated aplastic anemia mice. (**C-D**) Quantification of the absolute number (C), and frequency (D), of CD8+ and CD4+ lymphocytes. Frequency represented as percent indicated population among total bone marrow cells (BMCs). N=5-6 mice/group. Two-tailed student’s t-test used to compare Veh- versus PGDHi-treated mice for all parameters.

### PGDHi augments the effects of low-dose cyclosporine on cytopenias and BM cell loss

While IST is the front-line therapy for AA patients not receiving a bone marrow transplant, IST refractoriness is observed in some patients and IST dosing is often limited by neurological, hepatic, and renal toxicity (20). Given the immune-independent nature of PGDHi protection in our AA mouse model, we questioned whether PGDHi could increase the efficacy of sub-therapeutic low-dose cyclosporine A (CsA). We first compared responses to conventional-(50mg/kg) and low-dose (10mg/kg) CsA in our model. Conventional CsA dosing led to a complete protection of blood cell counts, whereas low-dose CsA-treated mice remained leukopenic, neutropenic, and thrombocytopenic (**Supp. Fig. 4A**). While conventional dosing prevented AA-related cytopenias, mice displayed signs of neuro- and hepato-toxicity as evidenced by hyperactivity and yellow pigmented serum with elevated total serum bilirubin (**Supp. Fig. 4B**).

Having identified low dose 10mg/kg CsA as a well-tolerated but subtherapeutic IST dose in our model, we tested the capacity for combination with PGDHi to confer an additive therapeutic effect. Coadministration of PGDHi and low-dose CsA significantly increased total white blood cells, neutrophils, platelets, and red blood cells (**Fig. 3A**) at 13 days post disease induction, with the combination increasing neutrophil levels significantly above those for monotherapy with either PGDHi or with low-dose CsA alone. Although dual therapy yields absolute neutrophils counts (mean 478 neutrophils (NE)/ul blood) just below the therapeutic threshold of PMN #500, these animals would not be considered at serious risk for infection (generally >200-400/uL) (5).

**Figure 3:**
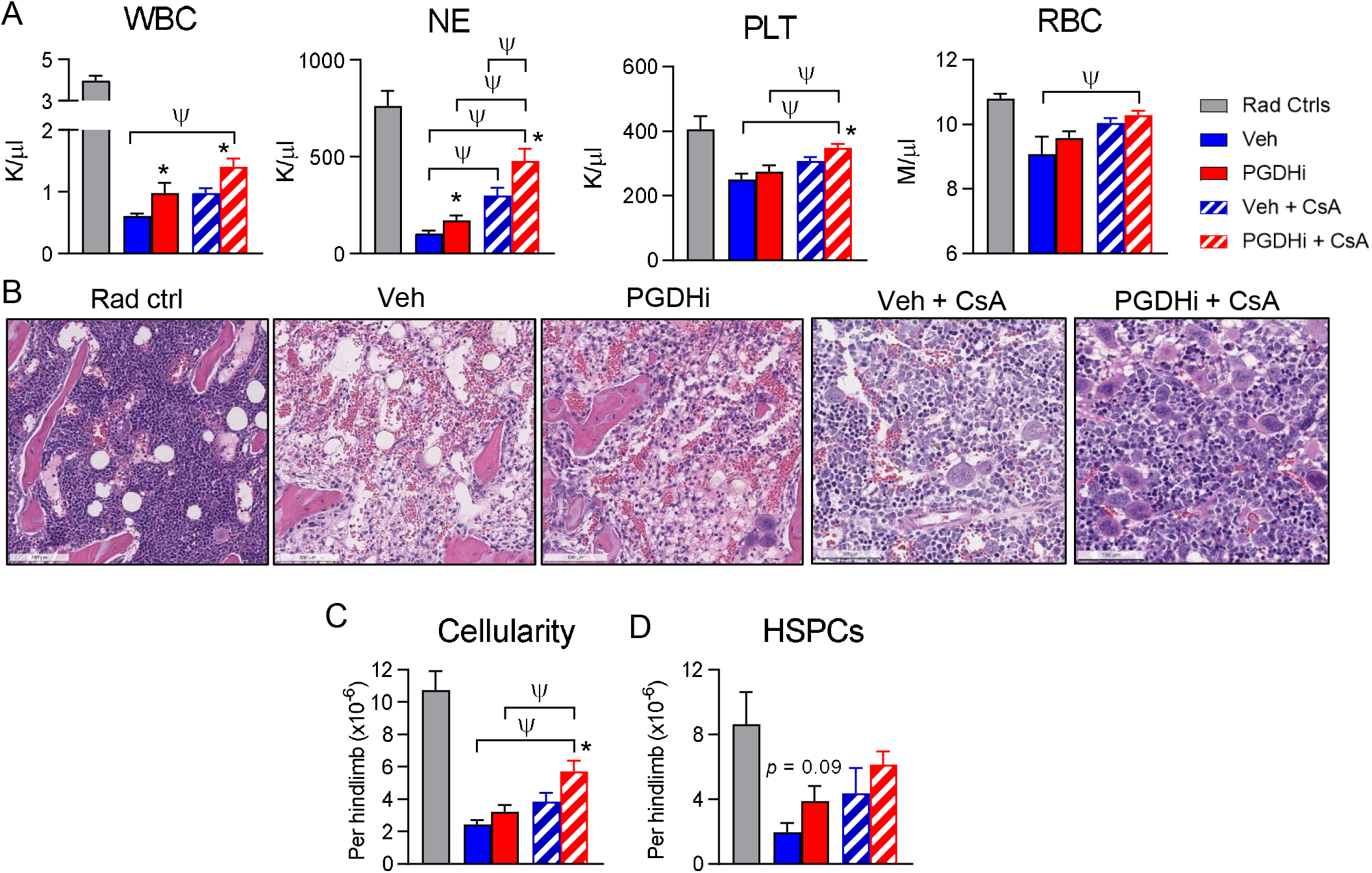
PGDHi improves the efficacy of low-dose cyclosporine in murine aplastic anemia. (**A**) Complete blood count analysis of radiation alone controls (Rad Ctrl), and aplastic anemia mice treated with PGDHi, vehicle (Veh) control, PGDHi + 10mg/kg cyclosporine (CsA), and Veh + CsA, 13 days post-induction. Panels represent total white blood cells (WBC), neutrophils (NE), platelets (PLT), and red blood cells (RBCs). N= 11-12 experimental mice/group. (**B**) Representative hematoxylin and eosin-stained sections from the hindlimbs of Rad Ctrl and aplastic anemia mice treated as indicated, 14 days post-induction. (**C-D**) Quantification of bone marrow (BM) cellularity and absolute number of BM Lin^−^ c-Kit^+^ -defined hematopoietic stem and progenitor cells (HSPCs) per hindlimb, 14 days post-induction. N=10-11 experimental mice/group. * P < 0.05 for the comparison of either mono- or dual-PGDHi therapy versus vehicle-treated counterpart by student’s t-test; ψ P < 0.05 by one-way ANOVA with Tukey multiple comparisons post-test.

Histologic evaluation of the BM at day 14 showed that concomitant with increasing peripheral blood counts, combination therapy also significantly preserved marrow architecture and morphologic hematopoiesis relative to vehicle-treated mice (**Fig. 3B**). In agreement with this observation, combination CsA + PGDHi significantly increased BM cellularity over that of mice treated with CsA alone. (**Fig. 3C**). Furthermore, CsA+PGDHi resulted in a trend towards increased numbers of phenotypic hematopoietic stem and progenitor cells (HSPCs; Lineage^−^ c-Kit^+^ cells; **Fig. 3D**), and phenotypically-defined stem cells (Lin^−^ c-Kit^+^ CD48^−^ CD150^+^ cells; **Supp. Fig. 5**), compared to CsA alone.

## Discussion

In the present study, we identify 15-PGDH as a novel therapeutic target that both independently protects hematologic function and complements front-line IST in a mouse model of aplastic anemia. In addition to IST, the c-Mpl agonist eltrombopag is frequently used to treat AA-related cytopenias (1, 2, 6). As PGDHi acts via an independent mechanism, we speculate it may also potentiate responses to eltrombopag. Importantly, AA is associated with clonal evolution of myeloid progenitor cells with the development of myelodysplastic syndrome and acute myeloid leukemia in 10-20% of cases (21). As we have previously established that PGDHi does not promote myeloma and leukemia xenograft growth (15), treatment with PGDHi should be a safe approach to improving therapy for AA. As, PGDHi also accelerates hematologic recovery and expands HSCs following transplant (15, 16), it may also have a role in improving recovery in AA patients receiving curative transplants. In overview, PGDHi monotherapy limits murine AA severity and is additive in improving therapeutic efficacy of low dose CsA. PGDHi may thus have potential to benefit long-term hematologic function of human AA patients and to enable treatment with better tolerated low-dose IST.

## Supplemental Methods

### Reagents

15-PGDH inhibitor (+)SW033291 was previously described (16), and provided by Dr. Sanford Markowitz (Case Western Reserve University, Cleveland, OH). (+)SW033291 was administered by intraperitoneal injection, twice per day spaced by 8 hours, beginning four days post-aplastic anemia induction. (+)SW033291 was prepared in a vehicle of 10% ethanol, 5% Cremophor EL, 85% dextrose-5 water, at a concentration of 125ug/200ul for use at 5mg/kg for a 25g mouse. Cyclosporine A (Sigma) was diluted in 10% ethanol, 5% Cremophor EL, and 85% IMDM to 250ug/200ul for use at 10mg/kg for a 25g mouse or to 1.25mg/200ul for use at 50mg/kg for a 25g mouse.

### Animals

Mouse experiments were conducted in the Case Western Reserve University Animal Resource Center, in accordance with guidelines of the Institutional Animal Care and Use Committee. Female C57BL/6J (H^b/b^) and CB6F1/J (H^b/d^) mice were obtained from Jackson Laboratories at 6 weeks of age. Mice were housed in standard microisolator cages and maintained on a defined, irradiated diet and autoclaved water. For aplastic anemia induction, mice were sublethally irradiated (3Gy) 4-6 hours prior to the retro-orbital infusion of 8e6 cells derived from the inguinal and axillary lymph nodes of age-matched C57BL/6J mice.

### Histological and immunohistochemical analysis

Animals were harvested via CO2 inhalation followed by cervical dislocation and hindlimbs were excised and placed in 10% neutral buffered formalin for 24 hours. Samples were transferred to PBS and shipped to Histowiz Inc. where they were embedded in paraffin, and sectioned at 4μm. Immunohistochemistry was performed on a Bond Rx autostainer (Leica Biosystems) with enzyme treatment using standard protocols. Bond Polymer Refine Detection (Leica Biosystems) was used according to manufacturer’s protocol. After staining, sections were dehydrated and film coverslipped using a TissueTek-Prisma and Coverslipper (Sakura). Whole slide scanning (40x) was performed on an Aperio AT2 (Leica Biosystems).

### ELISA

Peripheral blood from mice 14 days post-aplastic anemia induction was collected into Microtainer serum-separator tubes (Becton-Dickinson) by submandibular cheek puncture. Whole blood was allowed to clot at room temperature and then spun at 6000 x g for 3 minutes to separate serum. Serum was removed and stored at −80 prior to analyzing with the mouse IFNg Quantikine ELISA kit (R&D Systems), a custom Luminex (R&D Systems) for CXCL-12, G-CSF, and M-CSF, or the V-PLEX ProInflammatory Panel 1 Mouse Kit (Meso Scale Diagnostics) for all other analytes.

### Complete Blood Count Analysis

Peripheral blood from mice 13 days post-aplastic anemia induction was collected into Microtainer EDTA tubes (Becton-Dickinson) by submandibular cheek puncture. Blood counts were analyzed using a Hemavet 950 FS hematology analyzer. Values were tabulated graphically with error bars corresponding to standard error of the means and compared using 2-tailed t-tests.

### Flow Cytometry

Bone marrow was obtained by flushing the hindlimbs of radiation control and AA mice 14 days post-induction. Marrow cellularity was measured following red blood cell lysis. Cells were stained with the antibodies indicated in Table 1 and data was acquired on an LSRII flow cytometer (BD Biosciences). Immunophenotypic analysis was performed on FlowJo software (TreeStar).

### Colony Forming Analysis

1e5 total bone marrow cells were plated in Methocult media M3434 (StemCell Technologies) supplemented with hemin. CFU-GM and BFU-E colonies were scored 12 days post-plating.

### Statistical Analysis

Analysis was performed using GraphPad Prism software. Two-tailed Student t-test was used to compare between groups, unless otherwise noted. A Log-Rank (Mantel-Cox) test was used compare the overall survival time of mice that received C57BL/6J lymphocytes. Significance was established at P < 0.05.

## Acknowledgments

This work was funded by NIH grants R35 CA197442 and K99 HL135740, and was supported by the Radiation Resources Core Facility (P30CA043703), the Hematopoietic Biorepository and Cellular Therapy Core Facility (P30CA043703), the Tissue Resources Core Facility (P30CA043703), and the Cytometry & Imaging Microscopy Core Facility of the Case Comprehensive Cancer Center (P30CA043703).

## Supplemental Figure Legends

**Supplemental Figure 1: Red blood cell counts are not impacted by PGDHi.** Complete blood count analysis of red blood cells (RBCs) in radiation control, and aplastic anemia mice treated with PGDHi and vehicle control, 13 days post-induction. N=21 experimental mice/group.

**Supplemental Figure 2: PGDHi does not significantly impact circulating inflammatory factors.** Measurement of indicated factors in peripheral blood serum of mice, 14 days post-induction, in radiation control (Rad Ctrl), and aplastic anemia mice treated with PGDHi and vehicle control. N=4-18 mice/group. Two-tailed student’s t-test used to compare Veh- versus PGDHi-treated mice for all parameters.

**Supplemental Figure 3: PGDHi treatment does not reduce T cell expression of activation markers.** Mean fluorescence intensity (MFI) of CD90, Fas Ligand (FasL), and CD11b on CD8+ **(A)** and CD4+ **(B)** T lymphocytes. N=6 experimental mice/group. Two-tailed student’s t-test used to compare Veh- versus PGDHi-treated mice for all parameters.

**Supplemental Figure 4: 10mg/kg cyclosporine A is subtherapeutic but has better liver tolerability than standard dose.** (**A**) Complete blood count analysis of radiation alone control (Rad Ctrl), and aplastic anemia mice treated with low (10mg/kg), and conventional (50mg/kg) cyclosporine (CsA), 13 days post-induction. Panels represent total white blood cells (WBC), neutrophils (NE), and platelets (PLT). N= 6 experimental mice/group. (**B**) Measurement of total Bilirubin (TBILI) in peripheral blood serum of mice treated as indicated, 14 days post-induction. N=6 experimental mice/group.

**Supplemental Figure 5: PGDHi attenuates phenotypic HSC depletion**. Quantification of phenotypic hematopoietic stem cells (HSCs), defined as Lin^−^ c-Kit^+^ CD48^−^ CD150^+^ cells, in the bone marrow 14 days post-induction in radiation control (Rad Ctrl), and aplastic anemia mice treated as indicated. N= 10-12 experimental mice/group.

**Supplemental Figure 6: Dual administration of PGDHi and low-dose CsA does not impact serum IFN**γ. Measurement of IFNγ in peripheral blood serum of radiation control (Rad Ctrl), and aplastic anemia mice treated with Veh + CsA and PGDHi + CsA, at day 14. N=12 experimental mice/group.

