## Supplemental Table for "Inhibition of 15-PGDH Protects Mice from Immune-mediated Bone Marrow Failure"

Table 1. Flow cytometry antibodies used.

| Antibody | Clone | Manufacturer |
|---|---|---|
| CD45R/B220 | RA3-6B2 | BD Pharmingen |
| CD11b | M1/70 | BD Pharmingen |
| CD3e | 500A2 | BD Pharmingen |
| Ly-6G and Ly-6C | RB6-8C5 | BD Pharmingen |
| TER-119 | TER-119 | BD Pharmingen |
| Ly-6A/E (Sca-1) | D7 | Invitrogen |
| CD117 (c-Kit) | 2B8 | Invitrogen |
| CD48 | HM48-1 | eBioscience |
| CD150 | mShad150 | eBioscience |
| CD90 | 53-2.1 | eBioscience |
| CD8 | 53-6.7 | BioLegend |
| CD4 | RM4-4 | BioLegend |
| FasL | MFL3 | BioLegend |
| CD11a | M17/4 | BioLegend |
