## Supplemental Figures for "Inhibition of 15-PGDH Protects Mice from Immune-mediated Bone Marrow Failure"

S1. Red blood cell counts are not impacted by PGDHi monotherapy

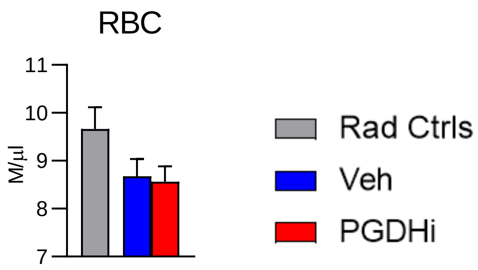

S2. PGDHi does not significantly impact inflammatory factors in the serum

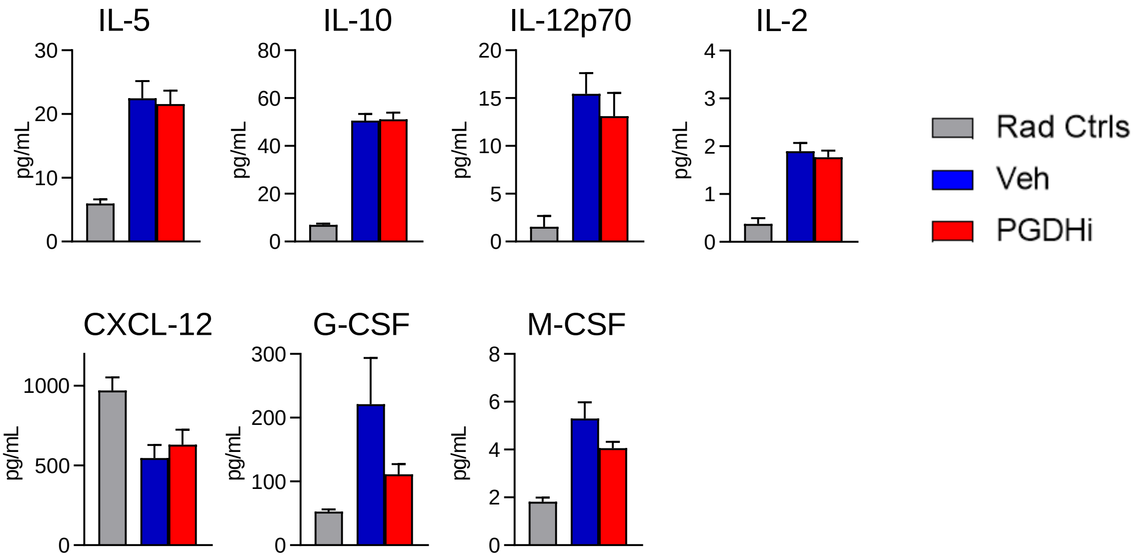

### S3. PGDHi treatment does not reduce T cell expression of activation markers

A

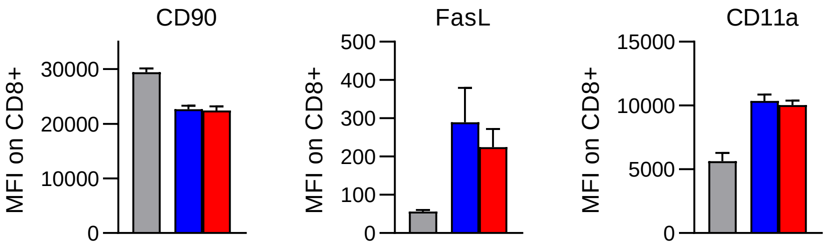


B

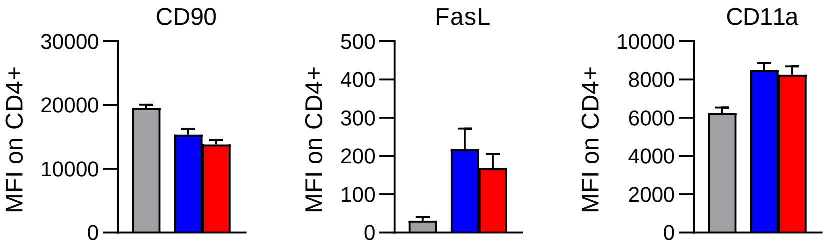

S4. 10mg/kg cyclosporine A is subtherapeutic but has better liver tolerability than standard dose

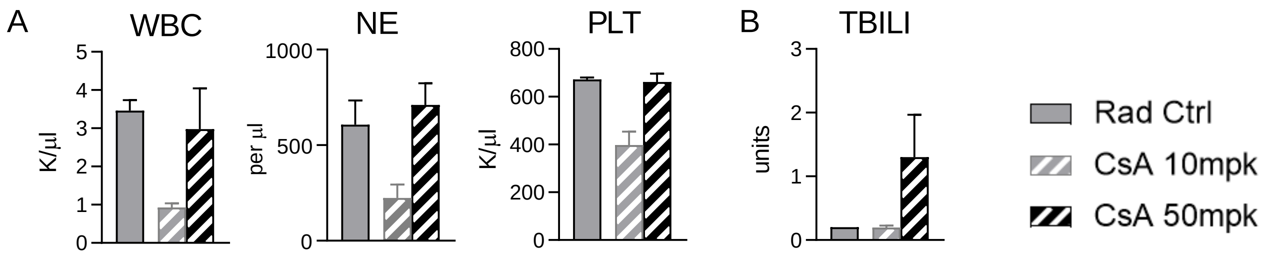

### S5. PGDHi attenuates phenotypic HSC depletion

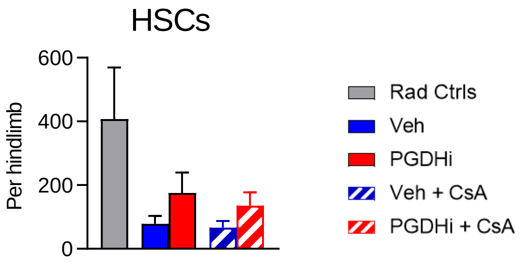

S6. Dual PGDHi and low dose CsA does not impact serum IFNγ

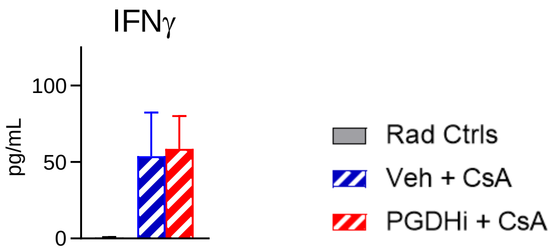
